# New sequential touch method for determining bacterial contact transfer rate from finger to surface

**DOI:** 10.1101/328971

**Authors:** Pengcheng Zhao, Yuguo Li

## Abstract

Bacteria can be transferred via surface touch. To evaluate the transfer rate, traditional single-touch methods require measuring the number of bacteria on donor and recipient surfaces, which is typically characterized by high levels of uncertainty. In this study, two concentrations of *Staphylococcus aureus* ATCC 25923 were inoculated on a clean thumb. For each set of trials, sequential touches were made between the thumb and each of 30 sterile glass slides, and each slide was placed in a sterile petri dish. The transferred bacteria on each slide were directly cultured in situ, and the colony-forming units (CFUs) were counted. The bacterial contact transfer rate was calculated by fitting the series of CFUs with the formula established. The average transfer rate was 12.9% under these conditions. The goodness of fit was compared in terms of the number of slides used in a set of trials and the number of CFUs counted on the slides. The use of more slides in a set of trials allowed more accurate evaluation of the transfer rate. The use of fewer than 20 slides was unacceptable. The high density of CFUs on the slides made counting them difficult, but if fewer than five CFUs were counted in a set of trials, the fit would be significantly influenced. To further evaluate the method, the dermal resident microflora on the thumb were also used to perform contact transfer tests. No statistically significant difference was found in the estimated transfer rate between the standard strain and the resident microflora.

**IMPORTANCE:** Diseases can be transferred indoors via the surface route because bacteria and viruses can be transferred to and from the hands when a fomite is touched. Various methods have been used to estimate the bacterial contact transfer rate between hands and surfaces. Evaluated transfer rates have had significant deviations and varied significantly across studies, partially due to the use of the single hand-surface touch method, inefficient hand/surface sampling, and complicated bacteria culture.

In this study, the bacterial contact transfer rate was evaluated with a new method involving sequential touches between a donor and a series of recipients. The bacteria on the recipients were cultured in situ without hand/surface sampling, which simplified the process of surface bacteria quantification. The new method significantly reduces experimental complexity, decreases random errors in the data, and provides a new method for understanding microbial transfers between surfaces.

## INTRODUCTION

Surface bacteria transfer can occur between human hands and surfaces in buildings by either direct or indirect contact (1-3). A high risk of cross-contamination occurs via the fomite surfaces in public places (4-7) and confined spaces (8, 9). Microorganisms are transferred to or from a solid surface when the surface is touched by humans (10-12). Some researchers have studied the spread of disease via the fomite surfaces by quantifying the transmission of bacteria or viruses between hands and various surfaces (13-15).

The concept of bacterial contact transfer rate is used to quantify the efficiency of bacterial transmission during contact between a hand and a surface; it is the ratio of the number of bacteria transferred to the recipient surface to the total number of bacteria on the donor surface within the contact area (16-19). In previous studies (20-25), the hand or the surface was first inoculated with the targeted bacterium to allow estimation of the transfer rate, followed by a single contact between the hand and surface and a series of procedures, including hand and surface sampling and bacterial incubation.

Bacterial transfer from a single touch inherently possesses a high degree of randomness (24). Many studies have calculated the transfer rate by using the measured number of bacteria on a touched finger and a surface (20-25). The results have included significant random errors because many factors can influence bacterial transfer during a single touch, including the surface type and condition (24, 26), transfer direction (27, 28), contact duration (29), residence time (18), and contact pressure (16), and many factors are difficult to control, such as surface wetness (27), impurities in the bacterial solution (30), and the specific gesture of touching (28). All of these factors can hardly be regulated in a single contact to allow more accurate evaluation of the transfer rate, with the exception of adding repeated tests, but at the cost of a heavier workload.

The surface bacteria sampling process also generates significant random errors (31). Various sampling methods have been applied to quantify the surface bacteria, including swabbing (22, 24), eluting with eluent (26, 32), and rubbing in a medium (33). Each has specific appropriate situations for use, but in general, these sampling methods do not have very high efficiency or reliability. For example, the most frequently used swabbing method can introduce significant errors by both absorbing and irrigating bacteria (31, 34).

In this study, a new method is presented to quantify the bacterial transfer from a finger to a solid surface. Instead of a single touch, each set of trials included sequential touches between a donor thumb inoculated with *Staphylococcus aureus* and each of 30 sterile glass slides. The transferred bacteria on each touched slide were *directly* cultured in situ without surface sampling, and the number of colony-forming units (CFUs) was counted. The transfer rate was obtained by fitting the series of CFU values with the established formula. Goodness of fit was compared in terms of the number of slides applied in a set of trials and in terms of the total CFUs counted on the series of slides. The thumb’s resident microflora were also used directly as the targeted bacteria for investigating the contact transfer rate.

## RESULTS

### Transfer of *S. aureus* from thumb to glass slide

A single touch was made between a donor thumb inoculated with *S. aureus* and each of 30 sterile glass slides in sequence as a set of trials. Six sets of parallel trials were performed with two magnitudes of bacterial concentration inoculated on the donor thumb (Figure 1, A-F). In all of the trial sets, the number of CFUs on the touched glass slides showed a decreasing trend as the sequence of contact progressed. The bacterial contact transfer rate was evaluated by fitting the numbers of CFUs counted on the series of touched glass slides with an established formula (Equation 4). The transfer rates were 11.4%, 12.8%, and 21.0% with the lower inoculated concentration and 11.7%, 12.1%, and 8.59% with the higher inoculated concentration. No statistically significant difference in transfer rate was seen between the two concentrations of bacteria inoculated on the thumb. The average transfer rate for the six sets of trials was 12.9%, with a small standard deviation (SD; 3.84%). The adjusted R-square values in the six sets of trials ranged from 0.371 to 0.809 (mean, 0.563), indicating a good data fit to evaluate the transfer rate.

**Figure 1.**
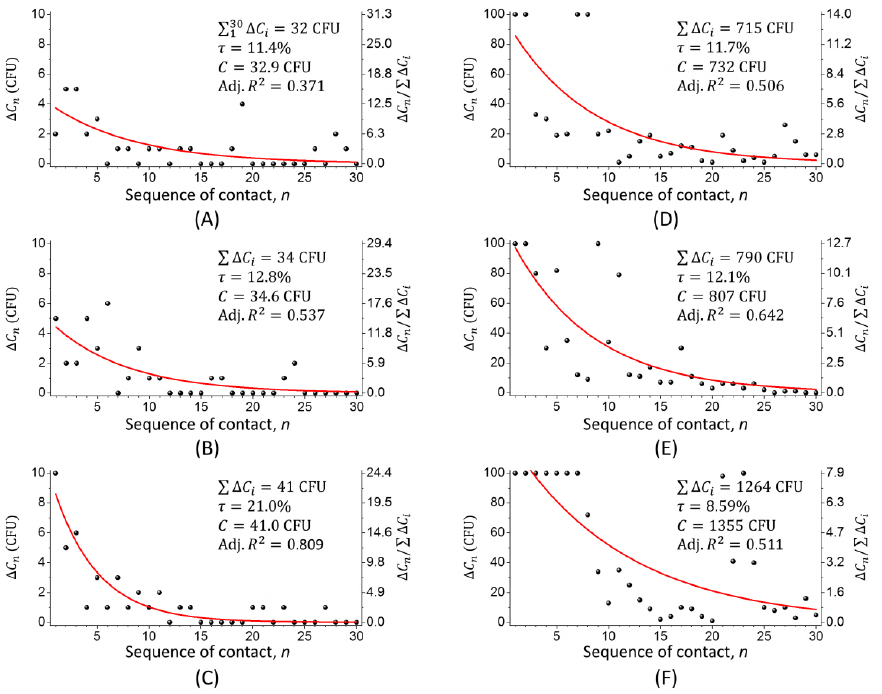
Transfer of *S. aureus* during sequential touches between a donor thumb and each of 30 sterile glass slides. Six sets of trials were performed with inoculated bacteria at a low concentration (A-C) and at a high concentration (D-F). For each trial, the CFUs counted on each slide are shown as the black spherical points, and the 30 CFU values fitted with Equation 4 are shown as the red curve. Based on the data fitting, the evaluated transfer rate, the evaluated initial number of bacteria on the donor thumb, and the adjusted R-square (Adj. *R*^2^) are listed for each set of trials.

### Different numbers of touches and CFUs

The reliability of the new method was analyzed in terms of the number of touches used in a set of trials and the number of CFUs counted on the touched glass slides. The evaluated transfer rate and adjusted R-square value are used as parameters to reflect the goodness of data fitting to allow comparisons between different numbers of touches and different numbers of CFUs (Figure 2).

**Figure 2.**
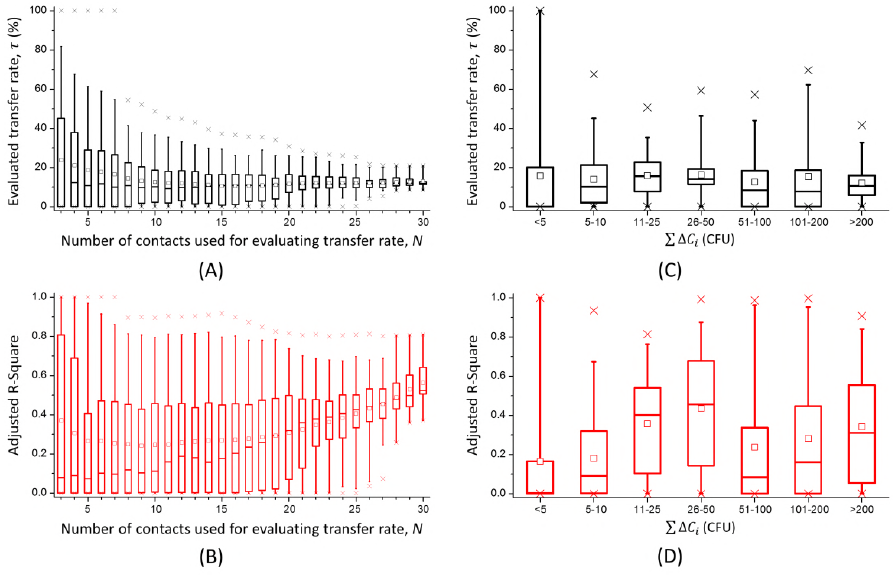
Evaluation of the goodness of data fitting in terms of the number of contacts and the total number of CFUs counted in a set of trials. Two parameters—the transfer rate and adjusted R-square—varied with the number of contacts (A and B) and the total number of CFUs (C and D), and were presented to evaluate the data fitting. The square, horizontal line, box, whisker, and cross labels in the box charts represent the mean, median, 25% to 75% range, 5% to 95% range, and 1% to 99% range, respectively.

Different amounts of data captured from the six arrays of 30 CFU values in Figure 1 were used for data fitting to evaluate the transfer rate. The use of more CFU values for data fitting, to represent a greater number of touches involved, resulted in smaller deviations in the evaluated transfer rate (Figure 2A) and larger adjusted R-square values in data fitting in each set of trials (Figure 2B). Significant deviations were seen in the evaluated transfer rate if fewer than 20 touches were made in a set of trials. Obvious decreases in deviations and significant increases in the adjusted R-square values were observed as the number of contacts was increased from 20 to 30.

The total number of CFUs counted for the series of touched slides had little influence on the transfer rate evaluation, but significant deviations in the evaluated transfer rate (Figure 2C) and unreliable data fitting in the evaluation (Figure 2D) were seen if fewer than five CFUs were counted for the whole series of touched slides.

### Transfer of dermal resident microflora

The contact transfer of skin bacteria was investigated. The resident microflora on the thumb were directly applied as the targeted bacteria for the contact transfer. A single contact was made between a donor thumb and each of 30 sterile glass slides in sequence as a set of trials. Six sets of parallel trials were performed, and the CFUs counted on the slides were fitted to evaluate the transfer rate (Figure 3). The average transfer rate was 12.8% (SD, 3.84%). No statistically significant difference in transfer rate was observed between the trials with inoculated *S. aureus* and those with the resident microflora on the thumb, but the adjusted R-square values for fitting the CFUs for skin bacteria transfer (mean, 0.432) were not as good as those with the model bacteria *S. aureus* (mean, 0.563).

**Figure 3.**
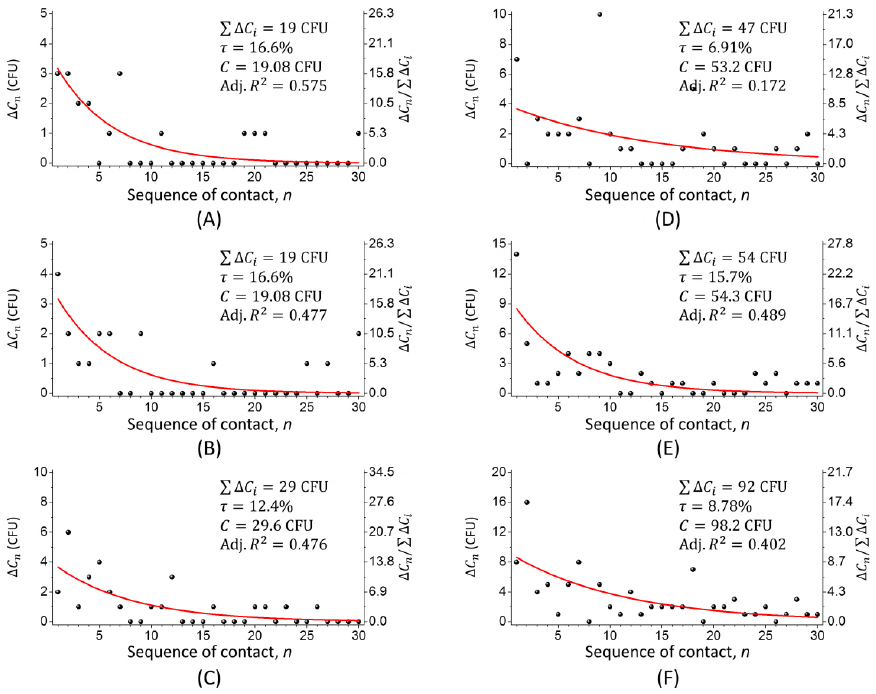
Transfer of dermal resident microflora on the thumb in sequential touches between a donor thumb and each of 30 sterile glass slides. Six sets of trials (A-F) were performed. For each trial, the CFUs counted on each slide are shown as the black spherical points, and the 30 CFU values fitted with Equation 4 are shown as the red curve. Based on the data fitting, the evaluated transfer rate, the evaluated initial number of bacteria on the donor thumb, and the adjusted R-square (Adj. *R* ^2^) are listed for each set of trials.

## DISCUSSION

The aim of this study was to propose a simple method to accurately evaluate the microbial contact transfer rate during surface contact. The process of surface bacteria quantification was simplified because the bacteria on the surface were cultured in situ with no surface sampling. Based on this simplification in bacterial culture, sequential touches can be performed between one donor and a series of recipients as a set of trials to evaluate one transfer rate value. Thus, a more accurate transfer rate was obtained with the sequential-touch method than with the single-touch method by fitting the CFUs on the series of recipients with the established formula.

In this new method to evaluate bacterial transfer from a thumb to a glass slide, a donor thumb inoculated with *S. aureus* was used to perform a set of sequential touches on a series of sterile glass slides. Therefore, the transfer rate was not evaluated from the CFUs on one contacted thumb and one glass slide, but from the CFUs counted on the entire series of recipient glass slides. The series of CFU values was fitted by Equation 4, introduced in Materials and Methods. The formula was established based on the assumption that the transferred bacterial amount decayed exponentially as the sequence of contact progressed. Figure 1 shows six sets of trials, with 30 contacts in each. In each set of trials, the number of CFUs on the touched glass slides showed a decreasing trend. Although the CFU values fluctuated significantly, the average adjusted R-square value of 0.563 indicates an acceptable level of data fitting.

For *S. aureus*, the average thumb to glass contact transfer rate was 12.9% (SD, 3.84%) in the six sets of trials with two magnitudes of inoculation concentration, as shown in Figure 1 (A-C, low concentration; D-F, high concentration). No significant difference was found in the transfer rates with different inoculation concentrations. It is believed that the bacterial contact transfer rate is low in dry conditions (24, 26, 35). A relatively low SD in the transfer rate values indicates a reliable evaluation from the new method. However, verification of this result against previous research is difficult because few studies have involved the transfer of bacteria from skin to nonporous surfaces. Also, a wide distribution of transfer rates has been presented by various studies with different experiment conditions, which makes comparison of our study and others difficult. Mackintosh and Hoffman (20) used several bacteria to study transfer from a finger to a porous surface (fabric) using a high inoculation volume, but they did not mention any process of surface drying. Only *Staphylococcus saprophyticus* had a comparable transfer rate (17%). For other bacteria, the transfer rates were much higher (88% for *Escherichia coli*, 76% for *Pseudomonas aeruginosa*, and 86% for *Klebsiella aerogenes*). Brar and Danyluk (26) studied the transfer rate of *Salmonella enterica* between gloved hands and tomatoes and measured transfer rates between 20% and 50% with a surface drying time of 0 or 1 hour and light pressure in contact. To the best of our knowledge, no studies have evaluated the bacterial transfer rate for contact from skin (naked finger without a glove) to nonporous surfaces. Lopez et al. (24) investigated the transfer rates between skin and glass surfaces for five types of microbes, but in the opposite direction; that is, the glass served not as the recipient but as the donor surface. They found the transfer rate of *S. aureus* to be 20.3% (SD, 33.4%) at a relative humidity between 15% and 32% and 45.5% (SD, 15.5%) at a relative humidity between 45% and 60%, but for *E. coli*, the transfer rates were 5.1% (SD, 5.4%) and 78.6 % (SD, 27.1%) under the two respective humidity conditions. In summary, the uncertainty of the surface wetness, the failure to note the physical parameters, and the lack of uniform sampling methods resulted in large variations and deviations in the evaluated transfer rates and made it difficult to generalize any trends from these studies as references. Therefore, few patterns in the evaluated transfer rate could be detected from these studies.

The parameters applied in the new method were analyzed. It is easy to see that a small deviation in evaluation of the transfer rate in this study was caused by sequential touches with a large number of glass slides and the significant number of CFUs counted on the slides. Figure 2A shows that the deviation in evaluating the transfer rate decreased as the number of touches applied increased, which means that the use of more recipient surfaces with the sequential-touch method can improve the accuracy of the transfer rate evaluation. Figure 2B shows that the average adjusted R-square value was greater than 0.3 when more than 20 touches were applied, and it continuously increased to 0.563 with 30 touches. Therefore, sequential touches with fewer than 20 contacts were unacceptable for the subsequent data fitting in this study. A larger number of contacts improved the accuracy with which the transfer rate was evaluated, but also increased the labor intensity. Therefore, a proper number of contacts should be determined while considering multiple factors when using the new method.

The bacterial concentration inoculated on a donor thumb should also be carefully controlled. Inoculation of a donor thumb with too high a bacterial concentration may result in difficulty in counting the CFUs on the touched glass slides, which is discussed below in detail. In contrast, the series of CFU values are meaningless and inappropriate for data fitting if the inoculated bacterial concentration is too low. Figure 2C shows a significant deviation in evaluation of the transfer rate if fewer than five CFUs are counted on the series of touched slides. In this case, more than 75% of the adjusted R-square values were smaller than 0.2, indicating meaningless data fitting with a group of small CFU values (Figure 2D). Therefore, a donor thumb inoculated with *S. aureus* at too low or too high a concentration results in an inaccurate evaluation of the transfer rate. In practical terms, fewer than five CFUs on a series of touched slides or more than hundreds of CFUs on each individual slide can affect the evaluation of the transfer rate.

The improvement in accuracy of the evaluation of bacterial contact transfer is mainly a result of the sequential-touch method used in the experiment. In the single-touch method, the eluate was subject to serial 10-fold dilutions and plate counts, yet only one or two CFU values from these plates fell within an expected range, which made it difficult to avoid random errors in calculating the transfer rate. The new method allowed us to determine the transfer rate by making sequential contacts in a set of trials rather than by a single contact while maintaining a low labor intensity. In sequential touches, a donor thumb makes a single touch on each recipient glass slide in sequence under constant conditions and parameters. Theoretically, the transfer rate remains constant during a set of sequential touches. The sequential touches reduced errors in evaluating the transfer rate by fitting the series of CFUs on the recipient with an ideal formula. The error in each calculated transfer rate was then neutralized by averaging the results from multiple touches.

The application of sequential touches is due to the simplification of the method of counting surface bacteria. In the traditional single-touch method, after a single touch, the bacteria on both the donor and recipient surfaces needed to be sampled. Swabbing the sampled surface followed by plate counting was the most widely used method for determining transfer rates (20-25). To sample bacteria, swabs were used to scrub various surfaces, but the efficiency and reliability of swabbing have both been shown to be very low and to vary across surfaces (31). In addition, the process of irrigating bacteria from the swab via vortex is often not well controlled, and other subsequent processes, such as sample dilution and transfer to a spread plate, often result in significant errors.

Some researchers have modified the sampling method via direct elution of bacteria from skin or other surfaces and then transferring the eluates of serial 10-fold dilutions onto spread plates for enumeration of the bacteria. The glove juice method was used to sample the bacteria from the hands (36). The sampling solution was poured into the worn glove from the wrist and sampled after a period of rubbing and irrigating. Conover and Gibson (37) followed the glove juice method to sample subjects’ hands when evaluating the efficacy of soaps. Lingaas and Fagernes (32) put the whole hand into a sterile bag instead of a glove for hand sampling and put the finger in a self-made finger stall for finger sampling. The European Norm (EN) 1500, as a standard method for evaluating the efficacy of hand hygiene, was used to quantify hand bacteria (38-40). Bellissimo-Rodrigues et al. (33) followed the EN 1500 standard to sample *E. coli* on fingers by directly rubbing the fingers in tryptic soy broth in a petri dish. Other methods have been attempted. Brar and Danyluk (26) put contaminated tomatoes and the removed glove into sampling bags with eluent and irrigated the bacteria into the eluent by shaking the bag. Sattar et al. (17) and Hübner et al. (41) eluted incubated bacteria by pressing a finger or fabric onto the mouth of the vial containing the eluent. This method is appropriate for sample bacteria from a specific area on skin or on some soft material surfaces. However, these modified methods are more labor-intensive and susceptible to manual operation (24), although the efficiency of surface bacterial sampling is improved. As with the EN 1500 method, the concentration of the irrigated bacteria in the eluate depended greatly on the volume of the five fingers merging into the eluent during sampling. Therefore, each method has a specific area of use, but none has been widely accepted as a replacement for swabbing in quantifying microbial transfer from surface contact (24).

In the method proposed in this study, bacteria were cultured in situ, which avoided the problems of surface sampling. Each sterilized glass slide used as a bacteria recipient was packed in a petri dish, as shown in Figure 4A. Immediately after contact by a donor thumb, the touched glass slides were covered directly with the warm culture medium (Figure 4B). After 4 days of incubation, it was easy to count the CFUs growing between the medium and glass slide (Figure 4C). In addition, because the bacteria were completely covered by the culture medium, the colonies were small and thin in the anoxic environment, which reduced the likelihood of the colonies mixing during culture. However, to ensure the number of CFUs would be well counted in each petri dish, an excessive concentration of *S. aureus* inoculated on the donor thumb had to be avoided. While the medium was poured, most of the bacteria remained immobile, although some flowed with the medium into the petri dish. Thus, most CFUs grew around the area at which the slide was touched by the donor thumb (Figure 4C). In this case, an excessive concentration of *S. aureus* on the donor thumb would have resulted in difficulty in counting the CFUs in the petri dishes with dense colonies grown on the touched slides. As shown in Figure 1, D-F, with a relatively higher concentration of *S. aureus* on the donor thumb, hundreds of CFUs were counted on some of the first few touched slides in each set of trials, which were difficult to count accurately. Therefore, the concentration of microorganisms used in this new method involving in situ culturing should not have been excessive.

**Figure 4.**
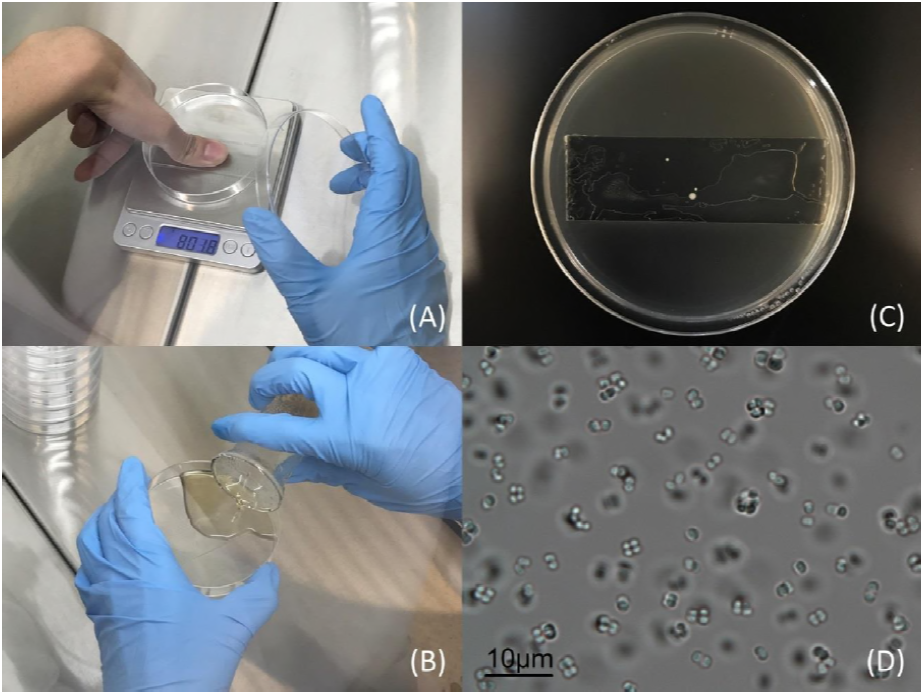
Test procedure of bacterial contact transfer from thumb to glass slides. (A) The inoculated thumb is used to press on a sterile glass slide inside a sterile petri dish. The pressure is controlled at 800 g by placing the slide and dish on an electronic scale. (B) Liquid warm PCA is poured into the petri dish, covering the touched glass slide. (C) The bacterial colonies after 4 days of growth under PCA. (D) A sample of bacteria from the dermal resident microflora on the thumb under an optical microscope.

The new method in this study was also applied to evaluate the contact transfer rate of the dermal resident microflora on the thumb. Based on Equation 4, both the contact transfer rate and the initial number of skin bacteria were evaluated. The bacteria found in the colonies incubated on surfaces included like Sarcina, Staphylococcus, Bacillus, and *Micrococcus tetragenus*. The average transfer rate of the resident microflora from the six sets of trials shown in Figure 3 was similar to that of *S. aureus*, as shown in Figure 1. No significant difference in the transfer rate was seen between the dermal resident microflora and *S. aureus* as one kind of transient microflora on the thumb. One possible explanation is that the bacteria applied in the trials might have had a similar property of adhering to skin and surface. Furthermore, the transfer rate might have been independent of the type of bacteria, regardless of whether it belonged to the dermal resident microflora or extraneous transient microflora. It is known that bacteria usually exist collectively and have a layer of adhesive coating outside their bacterial cells (42-44). In this case, the differences in contact transfer rates between types of microbes (20, 22, 24, 45) might have resulted from errors in the trials. This hypothesis should be verified further in subsequent studies.

In the proposed method, the number of bacteria on a donor surface can be evaluated by testing only recipient surfaces. The trials in Figures 1 and 3 present the evaluated number of bacteria on the donor thumb before any contact. The new method is applicable to situations in which direct testing of bacteria on the donor surface is difficult, such as with skin, porous surfaces, and food. Through contact with a series of sterile recipients and by developing a bacterial culture in situ, both the initial number of bacteria on the donor and the contact transfer rate can be calculated.

In some situations, bacterial quantities on both donor and recipient surfaces are difficult to evaluate by culturing in situ, for instance finger-finger (8, 35) or finger-clothing (20, 45) contact. In such scenarios, both surfaces have to be considered as donors, but using the method above, the initial number of bacteria on each donor can be estimated, and the contact transfer rate between them can be estimated by comparison.

Despite these potential applications, limitations also exist for culture in situ in our method. The initial number of bacteria on the donor surface must be controlled within a certain range. If hundreds of bacterial colonies grow on the test surfaces, the colonies are likely to combine, which may affect the CFU counts. However, the presence of too few bacteria growing on the recipient surfaces may result in large deviations when calculating the transfer rate.

## MATERIALS AND METHODS

### Materials

*S. aureus* ATCC 25923 and the dermal resident microflora on thumb were used in this study to investigate bacterial contact transfer. Figure 4D shows that the bacteria either existed separately or gather together as a larger group, with observed diameters ranging from 1 to 10 μm. Glass slides were chosen as the surface to receive the bacteria from the donor finger in the touch experiment. A sterile plastic petri dish was used to hold a sterile glass slide. Liquid lysogeny broth (LB) medium was used to culture *S. aureus*, and plate count agar (PCA) with LB medium was used to culture and count the bacteria left on each touched slide.

### Sterilization

Before the glass slides were touched by a donor thumb, they were sterilized. Each glass slide was first carefully washed and put into an autoclave for sterilization at above 121°C for 1 h. Each glass slide was then dried on an alcohol lamp and moved into a petri dish after cooling.

The PCA also needed to be sterilized before use. It was placed into a conical flask sealed by aluminum paper. To ensure the effectiveness of sterilization, the PCA was also sterilized at above 121°C for 1 h. The PCA was then put into a 45°C water bath to keep it warm for use.

### Handwashing

Hands should be washed before transfer experiments with *S. aureus* and with the dermal resident microflora on the thumb as the targeted bacteria. Hand skin is heavily colonized by bacteria, with a density of more than 10^4^ CFU/cm^2^ (46). To measure bacteria transfer effectively, the sample hands were washed before the experiment (47).

For handwashing in this study, participant’s hands were wetted and lathered with a commonly used hand soap. The hands were then scrubbed for 20 to 30 s and rinsed with clean running water. Finally, the hands were dried with an air dryer. The participant was instructed to avoid touching any surfaces with the sample finger before the touch experiment.

For the experiment with *S. aureus* contact transfer, the hands were repeatedly washed following the same procedure, but for the experiment with the resident microflora on the thumb, a single session of handwashing was sufficient. The remaining bacteria were found to be appropriate for observation of contact transfer. Because the resident microflora themselves are the targeted bacteria, the transfer of too few CFUs to the recipients can affect the statistical evaluation of the transfer rate. In addition, handwashing removes the body oils on the skin. If an unwashed finger presses on a glass slide, skin oil is likely to be transferred to the glass surface along with the bacteria, which would enable the bacteria to move around (42-44). This movement may blur the edges of the colonies and make them uncountable on the plate.

### Bacterial inoculation

For the transfer of *S. aureus*, the external standard bacterial strain should be inoculated on the thumb as a donor. *S. aureus* suspension with a concentration of 10^5^ to 10^6^ CFU/ml via 12-h culture in liquid LB medium at 35°C was used for surface incubation. The participant’s washed left thumb was incubated by spreading a 2-μl suspension of no dilution (or a 10-fold dilution) of the bacteria donor as the high (or low) concentration. This means that the donor thumb carried 200 to 2000 CFU in the high concentration and 20 to 200 CFU in the low concentration. The incubated thumb was allowed to visibly dry before contact began.

### Contact between thumb and glass slides

As shown in Figure 4A, each hand-surface contact experiment was conducted in a biosafety cabinet. With a single touch, the donor thumb was pressed against a sterile glass slide surface. The petri dish with the targeted slide was placed on an electronic scale to control the force of thumb pressure at 800 g. When pressure was loaded to this value, the thumb was held against the glass surface for 10 s (16, 22, 24, 28, 48, 49) while maintaining force within an acceptable range of 5%. If the pressure fell outside the acceptable pressure range, all of the samples in the sequential touch trial were discarded. At this pressure, the contact area between the thumb and glass surface was measured as 3.6 cm^2^. Therefore, the pressure was around 0.2 to 0.3 kg/cm^2^, which is consistent with values from existing studies (13, 21, 22, 28, 32, 35, 50).

A single touch was made between the donor thumb and 30 sterile glass slides in sequence as a set of sequential-touch trials. Two groups of studies were conducted. First, the thumb inoculated with *S. aureus* was used to perform six sets of trials. Two magnitudes of bacterial concentration were each used for three sets of trials. Second, the thumb with the dermal resident microflora was used as the donor for six sets of trials. In the second group of studies, to keep the CFU value sufficient for observation and enumeration of bacteria, the participant was allowed to perform only one experiment (one trial) per day. The temperature in the laboratory was maintained at 22°C ± 1°C and the relative humidity at 65% to 75%.

### Bacteria incubation and quantification

The CFUs on each glass slide after incubation of surface bacteria were counted as the number of bacteria transferred during the contact between the donor thumb and recipient glass slide. For incubation, the PCA was kept at 45°C in a water bath for preparation. After the glass slides were touched by the donor finger, they were covered with liquid PCA, as shown in Figure 4B, within 10 min. After the PCA solidified, the petri dish was inverted and stored in an incubator at 35°C for bacterial culture. The cultures were required to last longer than 4 days. For quantification of bacteria, the glass slide in each petri dish was checked for new colony growth every 24 h. Because the glass slides were covered with the LB medium, the bacteria were deprived of oxygen. In these anoxic environments, the colonies were small and thin and grew slowly, as shown in Figure 4C. Nevertheless, most of the colonies emerged within 2 days and gradually became thick and noticeable.

### Calculation of the bacterial contact transfer rate

#### Bacterial contact transfer rate

For surface contact, the transfer rate is used to quantify the efficiency of bacterial transfer from the donor surface to the recipient surface. During contact between a donor thumb and a sterile glass slide, not all of the bacteria are in real contact, and not all are removed from the thumb (44). This means there is a proportionality to the bacterial transfer, which is called the bacterial contact transfer rate. This is defined as the ratio of bacteria transferred to the total bacteria present within the contact area between the two surfaces, as shown in Equation 1.

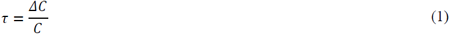

where the transfer rate in a single touch, *τ*, is equal to the ratio of *ΔC* to *C*. *ΔC* is the quantity of bacteria transferred during the touch, and *C* is the initial number of bacteria on the thumb before the touch.

#### Bacteria transfer in the sequential touches in a set of trials

Equation 1 suggests that, after a single touch between a donor thumb and a sterile glass slide, the number of bacteria left on the donor thumb is *C*-*ΔC = C-τC =* (*1-τ*)*C*. Assuming that a thumb is used to make a single touch with each of *N* sterile slides sequentially, the number of bacteria transferred to each slide is set as *ΔC*_*1*_, *ΔC*_*2*_, …, *ΔC*_*N*_. Then, in the sequential touches, the number of bacteria on the donor thumb will continue to decrease. Assuming that this series of contact actions is controlled under the same physical parameters, and that each bacterial particle’s probability of being touched and transferred is constant, the number of bacteria that remain on the donor thumb can be derived as below.

After the first touch, the number of bacteria left on the donor thumb is *C-ΔC*_*1*_ = (1- *τ*)*C*. After the second touch, it becomes *C-ΔC*_*1*_*-ΔC*_*2*_*=C-τC-τ*[(1*-τ*)*C*] = (1*-τ*)^2^*C*. After *N* touches, the bacteria quantity on the donor thumb can be obtained as *C-ΔC*_*1*_*-ΔC*_*2*_-…-*ΔC*_*N*_ = (1-*τ*)^*N*^*C*. A revised form is shown in Equation 2.

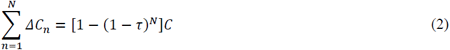

where 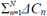 is the total number of bacteria transferred to *N* recipients over *N* touches. Equation 2 shows that as the number of touches (*N*) approaches infinity, the total number of transferred bacteria approaches the initial number of bacteria on the donor surface (*C*).

#### Bacteria transfer in each contact in a set of trials

According to Equation 2, the number of transferred bacteria (*ΔC*_*n*_, 1*≤n≤N*) during the *n*th touch in a set of trials can be derived as shown in Equation 3:

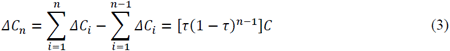

A relationship has then been constructed to connect the three parameters: the initial number of bacteria on the donor thumb (*C*), the bacterial transfer rate (*τ*), and the number of transferred bacteria in each touch in a set of trials (*ΔC* _*1*_, *ΔC* _*2*_, …, *ΔC* _*N*_). The number of transferred bacteria in each touch, the left part in Equation 3, will exponentially decay, which is the same mechanism as in radioactivity (51) and heat and mass transfer (52).

Ideally, the transfer rate *τ* can be directly obtained by *ΔC* and *C*, as shown in Equation 1. However, the initial number of bacteria on the donor thumb (*C*) is often unknown or difficult to measure. Following Equation 3, the donor thumb can be used to make identical touches on a pair of glass slides to obtain two sets of solutions of (*n, ΔC* _*n*_), that is, (1, *ΔC* _*1*_) and (2, *ΔC* _*2*_). With these observations, *τ* and *C* can be solved simultaneously. However, due to uncertainties in the measurements, using the two datasets also introduces significant uncertainty in the calculated transfer rate.

In the method of culture in situ that significantly simplified the experiment, sequential touches were made between a donor thumb and a series of sterile glass slides. An array of *ΔC* (*ΔC*_*1*_, *ΔC*_*2*_, …, *ΔC*_*N*_) was obtained by counting the CFUs on each subsequently touched slide. The values of *τ* and *C* in Equation 3 could then be evaluated more accurately by fitting the data with the formula introduced below.

#### Method for evaluating τ and C

Combined with Equation 2, the parameter *C* is removed from Equation 3 to reconstruct the relationship between the transfer rate (*τ*) and the number of transferred bacteria in each touch (*ΔC*_*n*_) in a set of trials, *ΔC*_*n*_ = ***f***_*τ*_(*n*), now with only one coefficient, *τ*, as shown in Equation 4. Using the least-squares method, the optimal solution for the transfer rate *τ* can then be determined. As such, the initial number of bacteria on the donor thumb (*C*) can be calculated using Equation 2.

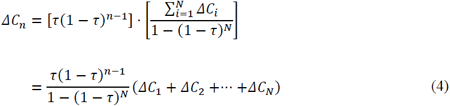

Equation 4 suggests that the transfer rate *τ* can be calculated indirectly by fitting the series of CFU values obtained from the first to the *N*th touches, with each counted as *ΔC*_*n*_(1 ≤ *n* ≤ *N*). This is the basis of the new sequential-touch method. Note that this method does not require evaluating the initial number of bacteria on the donor surface. This is replaced by a series of touches on clean recipients, resulting in a sequence of data that enables us to use data fitting to obtain an optimal value for the transfer rate.

### Statistical analyses

Origin 8.6 was used for data analysis and figure drawing. Student’s *t* test was performed to determine whether a statistically significant difference was found in the transfer rate when using different concentrations of inoculated *S. aureus* and when using *S. aureus* and the dermal resident microflora as the targeted bacteria for the experiment.

## ACKNOWLEDGEMENTS

This project is supported by a HKSAR Government CRF project (no. C7025-16G) and a GRF project (no. 17249616).

